# A Functional Resting-State Network Atlas Based on 420 Older Adults with Hypertension

**DOI:** 10.1101/2025.11.26.690831

**Authors:** Norman Scheel, Zac Fernandez, Josh Baker, Pavel Yanev, Jeffrey N. Keller, Ellen F. Binder, Eric D. Vidoni, Jeffrey M. Burns, Ann M. Stowe, Diana R. Kerwin, C. Munro Cullum, Linda S. Hynan, Wanpen Vongpatanasin, Rong Zhang, David C. Zhu

## Abstract

The *Risk Reduction for Alzheimer’s Disease* (rrAD) trial included 513 cognitively normal, sedentary, hypertensive older adults (aged 60 to 85 years) with dementia risk factors. We utilized 420 high-quality baseline resting-state functional MRI (rs-fMRI) scans from this cohort to develop a functional atlas tailored for aging populations. Typical rs-fMRI atlases derived from healthy young adults do not account for age-related changes, such as cortical atrophy, enlarged ventricles, and altered connectivity. To address this gap, we created a cohort-specific MNI-adjacent anatomical template, rrAD420, using SPM12’s DARTEL registration. In this space, we derived a comprehensive functional atlas using both group independent component analysis (GICA) and probabilistic functional mode decomposition (PROFUMO). The rrAD420 atlas offers detailed representations of Resting-State Network (RSN) connectivity, encompassing unique configurations and overlapping interactions. It features two Default-Mode Network (DMN)-specific seed-based maps (DMN24 with cerebellum, DMN18 without) and data-driven components resembling the major RSNs. Furthermore, PROFUMO allowed for the identification of multimodal and combinatory networks, capturing connections within and between RSNs. While optimized for hypertensive older adults, the rrAD420 atlas serves as a versatile tool for broader aging populations, aiding in the study of neurodegenerative processes and biomarker discovery.

## Introduction

Functional magnetic resonance imaging (fMRI) offers a noninvasive measurement of hemodynamic changes in the blood oxygenation level-dependent (BOLD) response to infer brain activity (Ogawa et al. 1990). Traditionally, fMRI studies utilize stimuli to investigate task-evoked BOLD signals to understand regional neuronal activation. Resting-state fMRI (rs-fMRI) offers a task-free approach that does not require a specific stimulus, examining spontaneous low-frequency (approximately 0.01 - 0.1 Hz) fluctuations in BOLD signals during rest conditions(B. Biswal et al. 1995). These BOLD signal fluctuations at rest preserve the synchronized oscillations of functionally connected brain regions observed during task-based fMRI (B. B. Biswal et al. 1995). Spatially distinct brain regions with highly correlated BOLD signal time courses are considered functionally connected and grouped into the same resting-state network (RSN). RSNs exhibit a consistent organization, and a comprehensive parcellation of RSNs forms a functional atlas (Damoiseaux et al. 2006; Fox and Raichle 2007; Shirer et al. 2012; Yeo et al. 2011). Functional atlases are widely used as a framework for understanding brain functional organization across individuals and populations.

Typically, RSNs are derived through seed-based or data-driven approaches, or by using existing atlases matched to the demographics of the population of interest for a given study. Seed-based approaches are considered hypothesis-driven, as the experimenter must choose an optimal seed region to generate the corresponding RSN map of interest (B. B. Biswal et al. 1995; Lalousis et al. 2022; Lowe et al. 1998). Seed-based approaches estimate the spatial map of regions sharing similar spontaneous BOLD signal fluctuations by cross-correlating the BOLD time course of voxels within the seed region with all other brain voxels’ time courses. In contrast, data-driven methods are exploratory, as they summarize the entire dataset into a fixed number of components or modes. These components or modes then provide multiple spatial configurations that resemble RSNs (Beckmann and Smith 2004; Damoiseaux et al. 2006; Farahibozorg et al. 2021; Harrison et al. 2015). Independent component analysis (ICA) is regarded as the current standard for data-driven decomposition of rs-fMRI data into distinct components with maximal spatial independence (Beckmann and Smith 2004). Recently, probabilistic functional modes (PROFUMO) emerged as another data-driven method for extracting RSNs in groups and individuals, without requiring spatial independence (Farahibozorg et al. 2021; Harrison et al. 2015). Regardless of the strategy used to characterize RSNs, the fundamental principle remains the same: brain regions grouped within the same RSN exhibit greater functional connectivity than those belonging to different RSNs.

Currently, there is no consensus on the exact number of distinguishable RSNs present in the brain, and RSN configuration can differ between functional atlases depending on the underlying population or the strategies used to define the RSNs (Doucet et al. 2019). However, there is a general consistency in the organization of major RSNs, such as the default mode network (DMN), executive control network (ECN), salience network, visual network, and somato-motor network (SMN) (Binder et al. 1999; B. B. Biswal et al. 1995; Cole et al. 2013; Cramer et al. 1999; Fox et al. 2005; Fox and Raichle 2007; Seeley et al. 2007; Shinn et al. 2015; Smith et al. 2009). Variability persists among functional atlases due to differences in demographics and RSN construction methodologies. Importantly, the demographic characteristics of individuals used to create functional atlases can influence RSN spatial configurations due to age-related neurobiological changes, including cortical atrophy, reduced white matter integrity, and altered functional connectivity patterns, which can reflect both a decline in network cohesion and/or compensatory mechanisms (Damoiseaux 2017; Doucet et al. 2019; Fjell and Walhovd 2010; Goh 2011; Madden et al. 2009; Zhu et al. 2010). Most existing functional atlases, such as those developed by Yeo (Yeo et al. 2011), Shirer (Shirer et al. 2012), and Damoiseaux (Damoiseaux et al. 2006), as well as anatomical standard image templates used for the spatial normalization for group-based results (e.g., MNI152, MNI305), are based on healthy young adults (Doucet et al. 2021; Evans et al. 1993; John Mazziotta et al. 2001; J. Mazziotta et al. 2001; Mazziotta et al. 1995). However, these templates do not adequately account for age-related structural and functional brain changes, posing challenges for their application in older populations (Damoiseaux 2017; Doucet et al. 2021; Fillmore et al. 2015; Madden et al. 2020).

Normal aging is associated with structural brain changes, including cortical thinning, gray matter atrophy, and ventricular enlargement, leading to substantial deviations from standard brain templates such as MNI152 and MNI305, derived from populations averaging 25.02±4.9 and 23.4±4.1 years of age, respectively (Evans et al. 1993; John Mazziotta et al. 2001; J. Mazziotta et al. 2001; Mazziotta et al. 1995; Raz 1997). These anatomical differences are problematic as they might cause severe artifacts when transforming non-normative brains into MNI space via volumetric non-linear normalization pipelines (Mazziotta et al. 1995). Functional connectivity also changes with age, even in the absence of pathological conditions (Damoiseaux et al. 2008; Doucet et al. 2021). For instance, Doucet et al. demonstrated that major RSNs, including the DMN, ECN, SMN, visual, and salience networks, exhibit different spatial patterns in older adults compared to younger adults (Doucet et al. 2021). Their findings emphasize the necessity for age-appropriate functional atlases and led to the development of Atlas55+, the first functional atlas derived from older adults. While the Atlas55+ provides an option to examine major RSNs in older adults, it lacks representations of common RSNs such as the language, limbic, and visuospatial networks (Shirer et al. 2012; Thomas Yeo et al. 2011).

To address these limitations, the present study aimed to develop a high-resolution, functional atlas in a standard space specifically tailored for older adults. We developed an age-appropriate standard anatomical space and a comprehensive RSN parcellation utilizing rs-fMRI and anatomical 3D T1-weighted image data from the *Risk Reduction for Alzheimer’s Disease* (rrAD) study. The rrAD study is a recently completed multisite randomized controlled trial designed to assess the effects of aerobic exercise and intensive cardiovascular risk factor management on neurocognitive function in hypertensive older adults with a family history of dementia and/or subjective cognitive decline (www.rradtrial.org) (Szabo-Reed et al. 2019; Szabo-Reed et al. 2023). The rrAD trial is registered at clinicaltrials.gov (NCT02913664). The comprehensive, harmonized neuroimaging protocols included anatomical, functional, and physiological MRI scans obtained at baseline and after 2 years of intervention. This new atlas is based on high-quality rrAD baseline MRI scans. It offers a robust framework for investigating brain functional connectivity in older populations, particularly those with hypertension and other age-related conditions.

One of the major contributions of this study is the development of a cohort-specific anatomical template, similar to the MNI template, termed *rrAD420*, from 420 high-quality baseline scans. In this space, we generated multiple functional parcellations using complementary methods: a hypothesis-driven seed-based analysis of the DMN; a group ICA (GICA)-based decomposition into spatially independent components (Beckmann and Smith 2004); and a PROFUMO-based decomposition allowing for overlapping and multimodal network configurations (Farahibozorg et al. 2021; Harrison et al. 2015). These approaches collectively offer a robust framework for studying the organization of RSNs in late adulthood.

The rrAD study provides a unique and robust dataset for examining the DMN and other RSNs in older adults. With its large sample size, standardized imaging protocols across multiple sites, and rigorous data quality assessments, the rrAD dataset is well-positioned to address the limitations of existing functional atlases for aging populations. The rrAD420 template and the corresponding functional atlas integrate various high-quality neuroimaging data into a shared anatomical space that accommodates age-related features such as cortical thinning and ventricular enlargement. By delivering anatomically and functionally relevant spatial maps derived from this population, the atlas enhances cross-study comparability. It supports investigations into the neural mechanisms of late-life cognitive decline, dementia prevention, and the identification of early biomarkers of neurodegenerative disease.

## Methods

### Participants

The rrAD sample cohort is comprised of 513 cognitively normal, sedentary hypertensive older adults with a family history of dementia and/or subjective memory complaints (Szabo-Reed et al. 2019). After quality assurance, baseline scans from 420 participants (age 60 to 85 years; 63% females; 34% aged 71-85; 13% African American; 4% Hispanic/Latino) were included in this study. Detailed baseline study participant characteristics are presented in Table 1 and in the corresponding publication (Szabo-Reed et al. 2023). The rs-fMRI and 3D T1-weighted images were collected across five different 3T MRI scanners. Each scan underwent rigorous manual quality control to ensure harmonization, as directed by a senior MRI physicist (Dr. Zhu). Participants were recruited from the Dallas, Baton Rouge, Kansas City, and St. Louis areas. This study was approved by the Human Subjects Committee at Pennington Biomedical Research Center (PBRC; n=128), The University of Texas Southwestern Medical Center (UTSW; GE scanner n=56; Phillips scanner n=44), University of Kansas Medical Center (KUMC; n=103), and Washington University School of Medicine (WashU; n=89). All participants met the strict inclusion criteria detailed in the rrAD rationale and methods paper and signed the necessary consent forms before participation (Szabo-Reed et al. 2019).

**Table 1:** Baseline characteristics of the full rrAD imaging cohort (N=434)**

| | 25% | Median | 75% | Mean ± SD | Minimum | Maximum |
| --- | --- | --- | --- | --- | --- | --- |
| Male/Female (n) |  |  |  | 160/274 |  |  |
| Race (n) |  | 360 Caucasian, 61 African American, 13 Hispanic |  |  |  |  |
| Age (years) | 64 | 68 | 72 | 69 ± 6 | 60 | 84 |
| Body mass index (kg/m<sup>2</sup>) | 26 | 30 | 34 | 30 ± 5 | 19 | 47 |
| Education (years) | 15 | 16 | 18 | 16 ± 2 | 12 | 20 |
| MMSE total | 28 | 29 | 30 | 29 ± 1 | 24 | 30 |
| Heart rate (bpm) | 59 | 68 | 75 | 68 ± 11 | 39 | 106 |
| Systolic blood pressure (mmHg) | 125 | 136 | 149 | 138 ± 16 | 103 | 188 |
| Diastolic blood pressure (mmHg) | 74 | 79 | 84 | 79 ± 7 | 56 | 116 |
| Cardiovascular risk score* | 9 | 14 | 24 | 18 ± 13 | 3 | 81 |

\*10-year Atherosclerotic Cardiovascular Disease (ASCVD) Risk based on AHA Pooled Cohort
\*\*Note: The final functional atlas (rrAD420) was derived from a quality-controlled subset of 420 participants from this cohort.

### Image Acquisition

This study employed five different 3T MRI scanners (a Philips Achieva and a GE Discovery MR 750W scanner at UTSW, a Siemens Skyra scanner at KUMC, a Siemens Prisma scanner at WashU, and a GE Discovery MR 750 W scanner at PBRC). To accommodate inter-scanner variability and harmonize data collected across sites, each scanner protocol was individually calibrated to ensure sequence harmonization. All scanners utilized a 32-channel head coil, except for the UTSW GE system, which used a 48-channel head-neck coil. For the present study, we focused on the rrAD baseline rs-fMRI and T1-weighted anatomical images. The rs-fMRI scans were acquired with a 2.5 s TR (time of repetition), a 28 ms TE (time of echo), and a 64 x 64 matrix with 3.4mm x 3.4mm pixels. A 3 mm slice thickness was used on all systems except the UTSW GE system, which used 3.4 mm slices to maintain a good signal-to-noise ratio with the 48-channel coil. Each rs-fMRI scan lasted 12 minutes, during which participants were asked to focus on a fixation cross. This ensures a consistent rest condition across participants, as RSNs are influenced by the participants’ condition during scan acquisition (Fernandez et al. 2022; Patriat et al. 2013). For all participants, the anatomical 3D 1-mm isotropic T1-weighted MPRAGE scans with cerebrospinal fluid suppressed were collected using the following parameters: 176 sagittal slices, TE = 3.8-4 ms, TR of acquisition ≈ 8.6 ms, time of inversion (TI) = 830 ms, TR of inversion = 2,330 ms, flip angle = 8°, FOV (field of view) = 25.6 cm x 25.6 cm, matrix size = 256 x 256, slice thickness = 1 mm.

### Data Preprocessing

For processing the rs-fMRI data in native space, we implemented an in-house routine based on the “afni_proc.py” routine from AFNI (Cox J.S. 1996), which generates a script for a standard preprocessing pipeline. For each participant, spikes in the signal time course were first detected and removed. Slice-timing corrections were applied to account for differences in acquisition time across slice locations. The functional images were co-registered to the T1-weighted high-resolution anatomical images, using the third volume as the reference. Then, we applied rigid-body motion correction in three translational and three rotational directions. Translational and rotational estimates, along with their derivatives, were modeled as regressors for the subsequent noise regression step. Spatial blurring with a full-width half-maximum of 4 mm was also applied to each fMRI dataset to reduce random noise and improve signal-to-noise ratio. Using the output from these initial preprocessing steps, we transformed the motion parameters from AFNI’s motion correction into an FSL-compatible format to perform aggressive ICA-AROMA to remove noise from participant motion from the rs-fMRI data (Pruim, Mennes, Buitelaar, et al. 2015; Pruim, Mennes, van Rooij, et al. 2015). As a final preprocessing step, we applied a temporal band-pass filter in the range of 0.009 Hz – 0.08 Hz as part of the regression model. A comprehensive explanation of our preprocessing steps can be found in our recent work, comparing different fMRI preprocessing strategies for older adults (Scheel et al. 2022).

After the rs-fMRI scans were co-registered to the T1-weighted high-resolution anatomical images, the origin within the T1 images was manually reoriented to the anterior commissure for each participant to stabilize the following spatial normalization steps. Then, to create a cohort-specific MNI-adjacent standard template, the reoriented T1-weighted images were used as inputs for SPM12’s segmentation and non-linear DARTEL registration (Ashburner 2007). We then transformed the rs-fMRI scans from the participants’ native space into the newly created standard rrAD420 standard space. The rrAD420 normalized anatomical space captures characteristics specific to the rrAD cohort and serves as the template for the functional connectivity profiles extracted from the rs-fMRI data (Figure 1).

**Figure 1:**
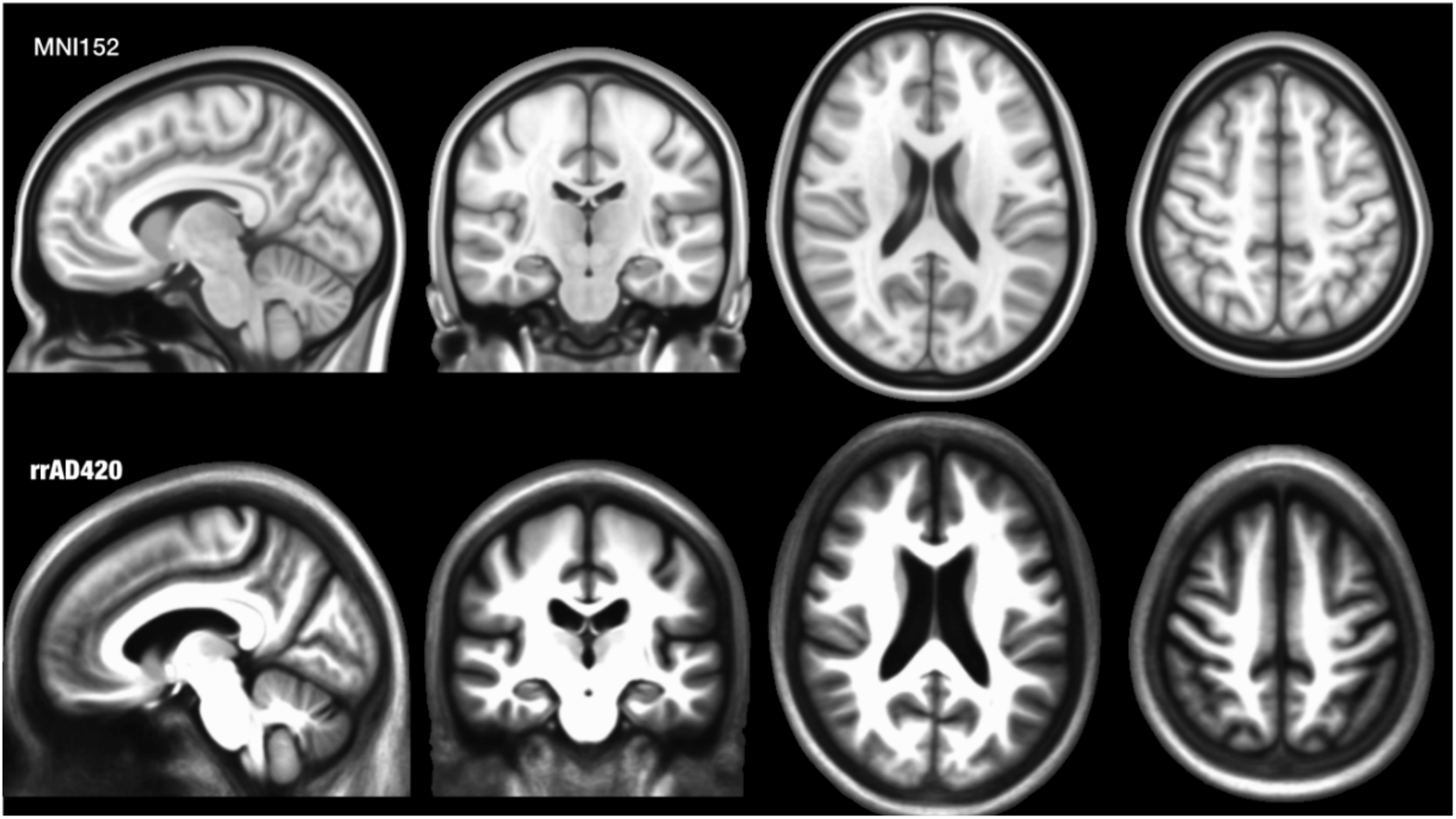
Comparison of standard MNI152 template (TOP) based on younger adults (25.02 ± 4.9) and the standard rrAD420 template based on 420 older adults (68.8 ± 5.9) in the rrAD study.

### Functional Data Decomposition

Using hypothesis-driven seed-based connectivity maps (B. B. Biswal et al. 1995; Cole et al. 2010; Lowe et al. 1998) in combination with data-driven approaches, temporally concatenated group independent component analysis (GICA) (Beckmann and Smith 2004) and probabilistic functional mode decomposition (PROFUMO) (Farahibozorg et al. 2021; Harrison et al. 2015), we created a functional atlas that describes the functional connectivity for older adults in the rrAD cohort. These methods offer complementary perspectives on network organization: a targeted, hypothesis-driven view of a single network (seed-based), a data-driven view of multiple, spatially independent networks (GICA), and a data-driven view of multiple, potentially overlapping networks (PROFUMO). These data-driven methods are developed for blind source separation (noise or signal) and allow for the identification of resting-state networks (RSNs) without requiring predefined hypotheses (Beckmann and Smith 2004; Cole et al. 2010; Harrison et al. 2015).

### Group Independent Component Analysis (GICA)

GICA is among the most widely used data-driven decomposition techniques in fMRI research. It is a matrix factorization method designed to unmix signals embedded in rs-fMRI data by identifying spatially distinct brain regions of interest (ROIs) with highly correlated BOLD time courses. These correlated regions are grouped into components, with each component representing an individual RSN. This approach is well-suited for identifying RSNs across individuals, as it aggregates data while preserving inter-individual variability through a dual regression approach (Beckmann and Smith 2004).

In this study, GICA was performed using FSL’s MELODIC software on the baseline rs-fMRI data from the rrAD cohort. Temporal concatenation of the rs-fMRI datasets across all participants enabled the extraction of group-level RSNs capturing overarching functional connectivity patterns while ensuring sensitivity to individual differences (Beckmann et al. 2009). Through this process, GICA decomposed the rs-fMRI data into group-level components as topological maps and corresponding time courses (Bijsterbosch et al. 2018; Hyvärinen and Oja 2000; McKeown and Sejnowski 1998). However, while MELODIC’s GICA identifies spatially independent components, it does not account for spatially overlapping functional networks (Bijsterbosch et al. 2019). This limitation can pose challenges for regions involved in multiple neuronal functions, as their participation in overlapping RSNs may not be fully captured.

### Probabilistic Functional Mode Decomposition (PROFUMO)

PROFUMO, a data-driven probabilistic approach, complements GICA by allowing spatially overlapping functional networks (Farahibozorg et al. 2021; Harrison et al. 2015). By leveraging Bayesian principles, PROFUMO models RSNs as probabilistic modes, providing a representation of functional connectivity that better aligns with the underlying physiology.

In contrast to GICA, PROFUMO employs a bidirectional approach to simultaneously model group-level and individual-level spatial maps. This is achieved through a top-down framework, in which group spatial maps normalize individual participant maps, and a bottom-up framework, in which participant maps iteratively update group maps. This iterative process continues until the group spatial maps align optimally with the participant spatial maps, capturing inter-individual variability more effectively (Harrison et al. 2020). PROFUMO performs Bayesian inference to optimize the estimated probability distributions of spatial and temporal components, considering both group-level and individual participant data (Harrison et al. 2015). The initial map in PROFUMO is generated by decomposing the group-averaged rs-fMRI data into spatially overlapping modes, providing a starting estimate for both group-level and individual-level spatial and temporal distributions.

A notable advantage of PROFUMO over GICA is that it does not assume spatial independence between modes (modes are analogous to components in GICA). This allows for spatial overlaps across modes, enabling multifunctional ROIs to be included in multiple modes. This capability is particularly valuable for regions involved in diverse neuronal functions. For this study, we used PROFUMO to unmix the preprocessed rs-fMRI data into distinct functional modes, each comprising spatially overlapping ROIs with highly correlated BOLD time courses. We utilized the full estimation of the original PROFUMO model. The original PROFUMO model for our cohort of 420 participants required approximately 3 TB of working memory for the concurrent estimation of group and individual-level modes. Each model computation took approximately one week using Michigan State University’s High-Performance Computing Center.

The combination of GICA and PROFUMO ensures a comprehensive characterization of RSNs, capturing both shared group-level connectivity patterns and individual-level variations. This dual approach enhances the robustness of the functional atlas and facilitates its application across a wide range of studies focusing on age-related and hypertensive populations.

### Atlas Creation

To construct a comprehensive functional atlas, we employed a multi-pronged methodological strategy, combining a hypothesis-driven approach with two distinct data-driven methods. First, a hypothesis-driven map of the DMN was created through traditional seed-based correlation analysis to provide a foundational benchmark for our cohort. For this analysis, we chose the isthmus of the cingulate cortex as our seed region due to its well-established role as a central hub of the DMN, its high resting-state metabolic activity, and its reliable connectivity with other core DMN regions(Zhu et al. 2013). We used the FreeSurfer segmentation output for each participant to identify and select this region as a seed, and then correlated the BOLD time course of this region with the time courses of every other voxel throughout the brain (Fischl 2012). We then transformed the connectivity maps from each participant into rrAD420 standard space using the previously calculated DARTEL transformation (Ashburner 2007). Finally, we obtained our group-level DMN spatial maps by averaging all participants’ connectivity maps, thresholding using Gaussian mixture modeling, and ROI extraction (Beckmann and Smith 2004).

While the data-driven techniques GICA and PROFUMO are powerful, accurately determining the number of true signal sources in a mixed signal is challenging, especially with high-dimensional data, as automated methods, such as those described by Minka (Minka 2000), often struggle to provide reliable estimates. While ICA can isolate some noise components, it’s challenging to consistently distinguish between neuronal signals and noise (Bhaganagarapu et al. 2013). Therefore, we performed multiple iterations of both MELODIC and PROFUMO to find the best-fitting model and parcellation for our functional atlas. Specifically, we ran seven instances of GICA with the dimensionality hyperparameter set at 21, 35, 49, 63, 77, 91, and 105, and three instances of PROFUMO with 30, 50, and 80 dimensions. We chose to start with a 21-component parcellation for GICA as suggested by previous literature and incrementally increased the number of components with each iteration (Beckmann and Smith 2004). We followed a similar reasoning when selecting the number of parcellations to use for PROFUMO. However, we had to limit the number of iterations due to the immense computational demand required to run PROFUMO. We then manually checked the output of each GICA and PROFUMO iteration separately and labeled each component as neuronal signal or noise following the ICA hand classification protocol presented by Griffanti et al. (Griffanti et al. 2017). Components and modes determined to be of neuronal origin were cross-referenced with probabilistic anatomical atlases and assigned a network designation based on their spatial overlap and similarity to established RSNs. To facilitate the labeling phase, we implemented an anatomical dual-regression approach, combining FreeSurfer segmentation and DARTEL normalization to create probabilistic representations of commonly used brain atlases, such as the automatic anatomical labeling (AAL) (Tzourio-Mazoyer et al. 2002), Brodmann (Rasser et al. 2004), Desikan (Desikan et al. 2006), Destrieux (Destrieux et al. 2010), Yeo (Thomas Yeo et al. 2011), and Shirer (Shirer et al. 2012) from MNI into rrAD420 space, allowing conversions between rrAD420 and other atlases. We used these atlases in rrAD420 space to cross-reference with our data and label components and modes as the RSN that best aligned through a manual three-rater system. This procedure required three trained neuroimaging researchers to agree on a network designation for each GICA component and PROFUMO mode. Once all components were appropriately designated as signal or noise, we used metrics of RSN representation, RSN splitting, and RSN grouping to select the parcellation dimension that optimally separated our data. RSN representation was decided based on the percentage of RSNs present relative to the 14 networks from the Shirer atlas (Shirer et al. 2012). RSN splitting was quantified by tallying the number of networks split over multiple components, while RSN grouping was quantified by the number of components that represented multiple RSNs. RSN splitting and RSN grouping address issues of overfitting and underfitting, respectively. If the dimensionality hyperparameter exceeds the actual number of RSNs in the data, overfitting occurs, and a single RSN can be split across multiple components or modes. Conversely, if the dimensionality hyperparameter is less than the actual number of RSNs in the data, underfitting occurs, and multiple RSNs will be improperly grouped into a single component or mode.

## Results

### Seed-Based Correlation Analysis

Here, we provide a focused, hypothesis-driven spatial map of the DMN generated through a traditional seed-based approach (Figure 2). This analysis resulted in a DMN spatial map comprising 24 ROIs: 18 cortical and subcortical, and six cerebellar. We call this DMN-specific atlas DMN24 (including cerebellum) or DMN18 (excluding the six cerebellar regions). Bilateral cortical and subcortical areas identified within the DMN24 spatial map include the posterior cingulum, precuneus/angular gyrus, medial frontal gyrus/medial orbital gyrus, superior frontal gyrus, anterior superior temporal sulcus, medial prefrontal thalamus, right hippocampal subiculum, parahippocampal place area, and inferior frontal gyrus (BA11). The cerebellar ROIs encompass the right medial cerebellum, cerebellar tonsil, and posterior cerebellum.

**Figure 2:**
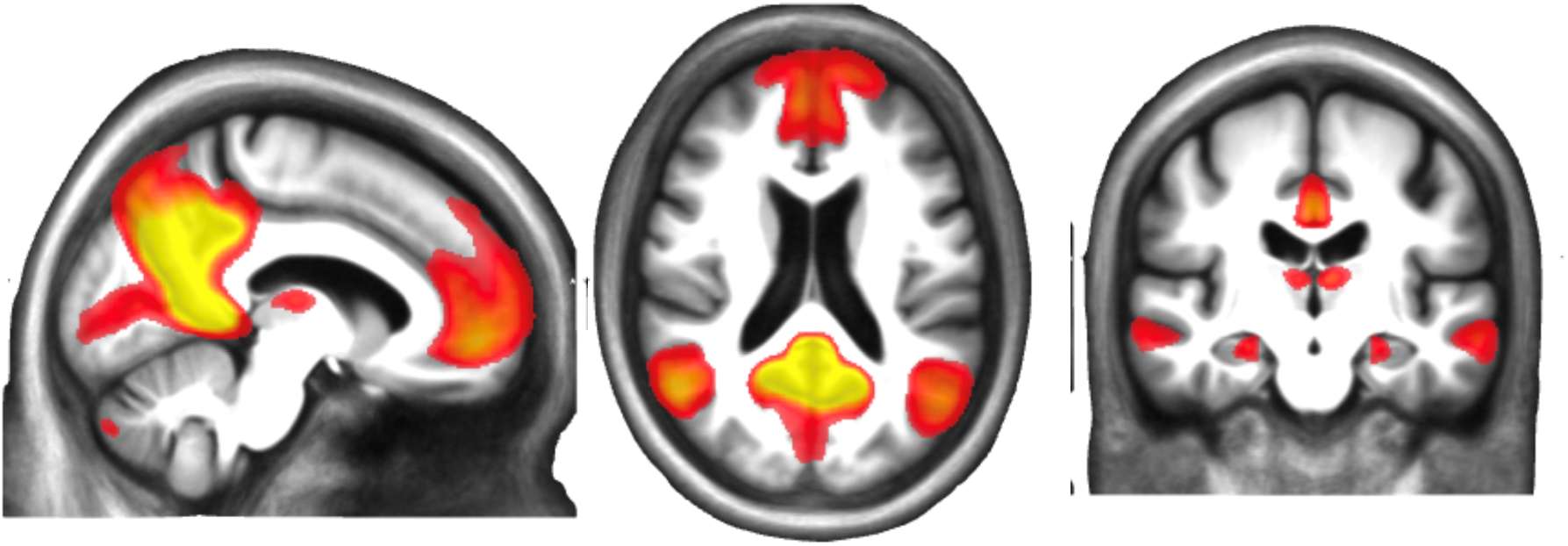
Representative images of the seed-based group spatial map of Default Mode Network (DMN24) based on 420 hypertensive older adults with a high familial risk of dementia or subjective cognitive decline that participated in the rrAD trial. The first 18 regions listed are cortical regions, while the remaining 6 are cerebellar regions.

### Group Independent Component Analysis

We examined the FSL MELODIC output for each of the seven GICA iterations, with the total number of components set to 21, 35, 49, 63, 77, 91, and 105 (Table 2). Adhering to the hand classification guide provided by Griffanti and colleagues, we categorized each component as either neuronal signal or noise (Griffanti et al. 2017). We found that the 21-component parcellation yielded nine components representing neuronal signals, while the remaining 12 were non-neuronal. Likewise, the 35-component parcellation yielded 15 neuronal-signal components and 20 noise components. The 49-component parcellation yielded 19 neuronal signal components and 30 noise components. While the remaining 63-, 77-, 91-, and 105-component parcellations yielded 20, 27, 31, and 32 components, respectively, that were likely derived from neuronal signals. We then designated each neuronal component as the RSN it best represented, using the 14 Shirer networks as a reference atlas (Shirer et al. 2012). In instances where a spatial map of a component represented the neuronal signal of an RSN that was not included in the Shirer atlas, such as the limbic network, we used additional functional atlases to supplement the labeling phase. To decide which parcellation strategy was best for our functional atlas, we assessed RSN representation, RSN splitting, and RSN grouping as defined in the methods section.

**Table 2:** FSL MELODIC output for the seven GICA iterations with the total number of components set to 21, 35, 49, 63, 77, 91, and 105, and the three PROFUMO iterations with the number of modes set to 30, 50, and 80. Following the hand-classification protocol outlined by Griffanti et al., each component was classified as either signal or noise. Compared to the 14-network reference atlas by Shirer et al. RSN representation describes the number of networks present as primary components. RSN splitting describes the number of components that split networks into multiple components. RSN grouping describes the number of components with multiple networks.

|  | Extracted components | Neuronal components |  | Noise components |  | Primary RSNs representation |  | RSN splitting |  | RSN grouping |  |
| --- | --- | --- | --- | --- | --- | --- | --- | --- | --- | --- | --- |
| **GICA** | 21 | 9 | 43% | 12 | 57% | 9 | 64% | 0 | 0% | 6 | 67% |
|  | 35 | 15 | 43% | 20 | 57% | 12 | 86% | 4 | 27% | 9 | 60% |
|  | 49 | 19 | 39% | 30 | 61% | 13 | 93% | 8 | 42% | 7 | 37% |
|  | 63 | 20 | 32% | 43 | 68% | 13 | 93% | 13 | 65% | 11 | 55% |
|  | 77 | 27 | 35% | 50 | 65% | 13 | 93% | 23 | 85% | 11 | 41% |
|  | 91 | 28 | 31% | 63 | 69% | 13 | 93% | 25 | 89% | 15 | 54% |
|  | 105 | 31 | 30% | 74 | 70% | 14 | 100% | 27 | 87% | 14 | 45% |
| **PFM** | 30 | 24 | 80% | 6 | 20% | 12 | 86% | 19 | 79% | 16 | 67% |
|  | 50 | 38 | 76% | 12 | 24% | 12 | 86% | 28 | 74% | 20 | 53% |
|  | 80 | 52 | 65% | 28 | 35% | 13 | 93% | 49 | 94% | 27 | 52% |

We found that all iterations yielded representations of all 14 Shirer networks in some capacity, either as a single network in a single component or as a secondary RSN in a component that combines more than one RSN. However, when considering only the primary RSN representation in components, we found variance across parcellations. Specifically, the 21-component and 35-component parcellations did not identify all networks as primary components. The 49-component parcellation captured a primary representation of most RSNs, and there was a plateau effect, with the 63-, 77-, and 91-component parcellations performing at the same level. The 105-component parcellation was the only iteration to show primary RSN representation for all RSNs in single components. The next metric we considered in our decision was RSN splitting. Although the 105-component parcellation captured all primary RSNs from the reference atlas, it also showed one of the highest rates of splitting single RSNs across multiple components. This issue was not as severe in other parcellations, as there was a trend that, for more components extracted, network splitting became more common. Lastly, we considered RSN grouping and found the inverse of the RSN splitting effects. The 21-and 35-component parcellations grouped more RSNs into a single component, thereby reducing the primary network representation in these iterations. Overall, we found the 49-component parcellation provided the best balance between network representation, splitting, and grouping. Therefore, we chose this parcellation solution as the basis for the GICA portion of our rrAD420 functional atlas.

Within the 49-component GICA parcellation, we identified fifteen recognizable RSNs, including primary visual, sensorimotor (SMN), left executive control (LECN), precuneus, dorsal attention/visuospatial (DAN/Vis), ventral default mode (vDMN), high visual, posterior salience, anterior salience, language, dorsal default mode (dDMN), right executive control (RECN), auditory, basal ganglia, and limbic networks. Following the methods outlined in Pruim et al., these RSNs were ordered based on spatial reproducibility, with the primary visual network being the most reproducible and the limbic network being the most difficult to reproduce (Pruim, Mennes, Buitelaar, et al. 2015; Pruim, Mennes, van Rooij, et al. 2015). A listing of the number and names of regions included for each RSN, along with a visual representation of each RSN’s spatial map, can be found in Table 3 and Figure 3.

**Figure 3:**
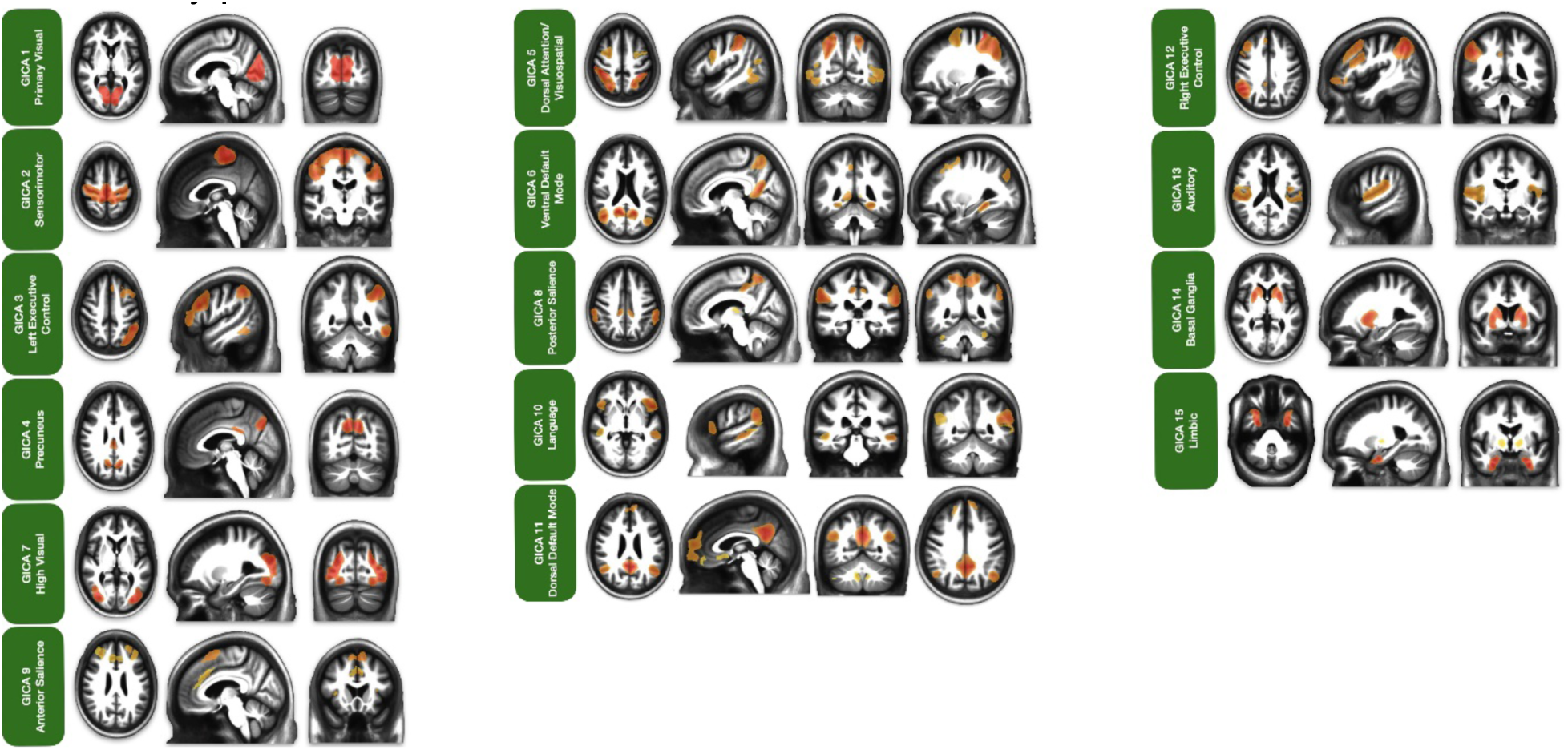
Representative spatial maps of RSN derived from 420 hypertensive older adults with a high familial risk of dementia or subjective cognitive decline, participants in the rrAD trial. Group independent component analysis (GICA) was performed on resting-state fMRI data using FSL’s MELODIC software. Displayed RSNs are from the 49-component parcellation, which provided optimal network separation among the tested models, capturing robust and well-defined functional connectivity patterns across the cohort.

**Table 3:** Complete RSN list from the consolidated 49-Component GICA Parcellation into 15 networks, including the number of ROIs comprising each RSN, and the anatomical locations of each of those ROIs.

| Network number | Network label | Number of ROIs | List of ROIs |
| --- | --- | --- | --- |
| 1 | Primary Visual Network | 2 | Superior middle occipital gyrus (striate cortex) |
| 2 | Sensorimotor Network | 4 | Superior pre/postcentral gyrus (Sensorimotor cortex), Inferior precentral gyrus (Sensorimotor cortex) |
| 3 | Left Executive Control Network | 6 | Left inferior parietal lobe/angular gyrus, Left middle/inferior frontal gyrus, Left superior medial gyrus, Left posterior middle cingulate cortex, Left inferior/middle temporal gyrus, Right posterior cerebellum |
| 4 | Precuneus Network | 4 | Precuneus, Posterior/middle cingulate cortex |
| 5 | Dorsal Attention/Visuospatial Network | 11 | Superior parietal cortex/precuneus, Superior frontal cortex, Inferior frontal gyrus, Middle temporal gyrus, Right inferior temporal gyrus |
| 6 | Ventral Default Mode Network | 10 | Precuneus, Frontal eye field, Middle occipital gyrus, Retrosplenial cortex, Parahippocampal place area |
| 7 | High Visual Network | 2 | Prestriate cortex |
| 8 | Posterior Salience Network | 10 | Supramarginal gyrus, Precuneus/middle cingulate cortex, Thalamus, Insula, Superior cerebellum |
| 9 | Anterior Salience Network | 14 | Superior frontal gyrus, Middle frontal gyrus, Dorsal anterior cingulate cortex; Superior frontal gyrus/pre-supplemental motor area, Insula, Superior cerebellum, Inferior cerebellum. |
| 10 | Language Network | 10 | Angular gyrus, Superior temporal gyrus, Superior/middle temporal gyrus, Pars triangularis, Middle temporal gyrus. |
| 11 | Dorsal Default Mode Network | 21 | Posterior cingulate cortex, Superior frontal gyrus Superior frontal/medial gyrus, Angular gyrus, Superior medial gyrus, Basal ganglia, Anterior cingulate cortex, Inferior frontal gyrus, Hippocampus, Middle temporal gyrus, Superior cerebellum, Inferior cerebellum |
| 12 | Right Executive Control Network | 8 | Right inferior parietal lobe/angular gyrus, Right middle/inferior frontal gyrus, Right superior medial gyrus, Right posterior middle cingulate cortex, Right pars opercularis/inferior frontal gyrus, Right middle frontal gyrus, Right inferior/middle temporal gyrus, Left posterior cerebellum. |
| 13 | Auditory Network | 2 | Superior temporal gyrus/operculum |
| 14 | Basal Ganglia Network | 2 | Basal ganglia |
| 15 | Limbic Network | 4 | Temporal pole, Pallidum |

### Probabilistic Functional Modes

To complement the GICA parcellation, we analyzed the outputs from the three PROFUMO iterations with the hyperparameter for the number of modes set to 30, 50, and 80 (Table 2). Using the same hand-classification protocol outlined by Griffanti et al. we categorized each mode as either neuronal signal or noise (Griffanti et al. 2017). The 30-mode parcellation yielded 24 modes of neuronal origin, while the remaining six modes were categorized as noise. The 50-mode parcellation identified 38 modes of neuronal origin and 12 noise modes, whereas the 80-mode parcellation included 52 modes of neuronal origin and 28 noise modes. We evaluated these parcellations using the same metrics of RSN representation, splitting, and grouping to determine the optimal parcellation for our functional atlas. In terms of RSN representation, we found that each iteration captured all RSNs of the reference functional atlas by Shirer et al.(Shirer et al. 2012). While not all networks were presented as primary components, the remaining networks emerged as combinatory networks. All in all, the 50-mode parcellation offered the best balance between minimizing RSN splitting (where a single network is fragmented across multiple modes) and reducing RSN grouping (where multiple distinct networks are inappropriately combined).

Within the 50-mode PROFUMO parcellation of the rrAD420 functional atlas, we identified 13 primary single RSNs, including core networks like the precuneus, primary visual, LECN, SMN, high visual, RECN, vDMN, dDMN, posterior salience, visuospatial, anterior salience, limbic, and basal ganglia networks. In addition, some modes integrated elements of multiple RSNs into a single mode. This is due to PROFUMO’s allowance for multimodal brain regions to be represented across multiple modes, whereas GICA seeks to maximize spatial independence for each region between components. These modes were kept in our functional atlas as “combinatory” networks. We identified six RSNs of this type, including the anterior ECN/dDMN (aECN/dDMN), language/dDMN, anterior salience/dorsal attention network (aSalience/DAN), language/temporoparietal, language/auditory/temporoparietal, and language/auditory networks. In contrast to the GICA portion of our functional atlas, the PROFUMO parcellation yielded cases in which multiple modes captured configurations of the same RSN. In these instances, all modes were kept as separate configurations for their respective primary RSNs. Nine RSNs of our functional atlas include sub-configurations, namely the primary visual, SMN, high visual, vDMN, dDMN, posterior salience, visuospatial, anterior salience, and limbic networks, as well as the language/DMN combined network. We assigned the mode that best captured the reference spatial map of its respective RSN as the primary representative configuration. As such, the primary representative network configuration was listed first for each RSN, with subsequent configurations in our functional atlas (i.e., “PROFUMO X.1” in Table 4).

**Table 4:** Complete RSN list from the 50-Mode PROFUMO parcellation, including the number of modes that comprise each network, the number of ROIs within each mode, and the anatomical locations of each of those ROIs.

| Network number | Network label | Subnetwork number | Number of ROIs | List of ROIs |
| --- | --- | --- | --- | --- |
| **SINGLE NETWORKS** |  |  |  |  |
| 1 | Precuneus Network |  | 6 | Precuneus, Posterior/middle cingulate cortex |
| 2 | Primary Visual Network | 2.1 | 2 | Superior middle occipital gyrus (striate) |
|  |  | 2.2 |  | Inferior occipital gyrus |
|  |  | 2.3 | 4 | Superior occipital gyrus, Middle occipital gyrus |
| 3 | Left Executive Control Network |  | 7 | Left inferior parietal lobe/angular gyrus, Left medial/inferior frontal cortex, Left superior medial frontal cortex, Left inferior/medial temporal cortex, Left posterior middle cingulate cortex, Right inferior semilunar lobule of the cerebellum, Right posterior lobule of the cerebellum. |
| 4 | Sensorimotor Network | 4.1 | 2 | Superior pre/postcentral gyrus |
|  |  | 4.2 | 2 | Inferior precentral gyrus |
|  |  | 4.3 | 2 | Middle precentral gyrus |
|  |  | 4.4 | 2 | Anterior cerebellum |
|  |  | 4.5 | 2 | Superior medial pre/postcentral gyrus |
| 5 | High Visual Network | 5.1 | 2 | Prestriate cortex |
|  |  | 5.2 | 2 | Inferior occipital gyrus |
|  |  | 5.3 | 2 | Superior occipital gyrus |
| 6 | Right Executive Control Network |  | 7 | Right inferior parietal lobe/angular gyrus, Right medial/inferior frontal cortex, Right superior medial frontal cortex, Right inferior/medial temporal cortex, Right posterior middle cingulate cortex, Left inferior posterior cerebellum, Left posterior cerebellum. |
| 7 | Ventral Default Mode Network | 7.1 | 10 | Precuneus, Frontal eye field, Middle occipital gyrus, Retrosplenial cortex, Parahippocampal place area |
|  |  | 7.2 | 2 | Precuneus |
| 8 | Dorsal Default Mode Network | 8.1 | 10 | Posterior cingulate cortex, Angular gyrus, Superior frontal gyrus, Medial prefrontal cortex, Middle temporal lobe |
|  |  | 8.2 | 2 | Posterior cerebellum |
|  |  | 8.3 | 2 | Prefrontal cortex |
|  |  | 8.4 | 4 | Anterior cingulate cortex, Middle cingulate cortex |
| 9 | Posterior Salience Network | 9.1 | 11 | Supramarginal gyrus, Precuneus/middle cingulate cortex, Superior middle frontal gyrus, Inferior frontal gyrus, Inferior middle frontal gyrus, Right middle temporal gyrus |
|  |  | 9.2 | 2 | Superior parietal lobule |
| 10 | Visuospatial Network | 10.1 | 8 | Superior parietal cortex/precuneus, Superior frontal cortex, Inferior frontal gyrus, Inferior temporal gyrus |
|  |  | 10.2 | 2 | Inferior parietal cortex and superior parietal cortex/precuneus |
|  |  | 10.3 | 3 | Right inferior parietal lobule, Right pars opercularis, Right middle frontal gyrus. |
|  |  | 10.4 | 2 | Inferior frontal gyrus |
|  |  | 10.5 | 2 | Intraparietal sulcus |
|  |  | 10.6 | 2 | Superior parietal lobule |
| 11 | Anterior Salience Network | 11.2 | 6 | Superior/middle frontal gyrus, Dorsal anterior cingulate cortex, Insula |
|  |  | 11.2 | 2 | Middle frontal gyrus |
|  |  | 11.3 | 2 | Secondary motor area (SMA) |
| 12 | Limbic Network | 12.1 | 2 | Orbitofrontal cortex |
|  |  | 12.2 | 2 | Temporal pole |
| 13 | Basal Ganglia Network |  | 2 | Basal ganglia |
| **COMBINATORY NETWORKS** |  |  |  |  |
| 14 | Anterior Executive Control Network/Dorsal Default Mode Network |  | 4 | Superior frontal sulcus, Medial superior frontal gyrus |
| 15 | Language Network/Dorsal Default Mode Network |  | 8 | Angular gyrus, Middle temporal gyrus, Left posterior cerebellum, Right pars triangularis, Right middle frontal gyrus, Right dorsomedial prefrontal cortex, Right precuneus. |
|  |  |  | 7 | Right posterior cerebellum, Left angular gyrus, Left middle temporal gyrus, Left pars triangularis, Left middle frontal gyrus, Left dorsomedial prefrontal cortex, Left precuneus. |
|  |  |  | 2 | Right pars triangularis, Left pars triangularis. |
| 16 | Anterior Salience Network/Dorsal Attention Network |  | 2 | Superior precentral gyrus |
| 17 | Language Network/Temporoparietal Network |  | 2 | Middle temporal gyrus |
| 18 | Language Network/Auditory Network/Temporoparietal Network |  | 2 | Posterior middle temporal gyrus |
| 19 | Language Network/Auditory Network |  | 2 | Superior temporal gyrus |

For the “single” networks, PROFUMO parcellation of the rrAD420 functional atlas demonstrated notable differences from the GICA-derived atlas, capturing distinct spatial and temporal features of RSNs (Table 4; Figure 4A). The precuneus network, represented in GICA with the posterior/middle cingulate cortex, was identified by PROFUMO as a single mode, isolating it more distinctly from other networks. This highlights differences in how the DMN’s core regions are represented.

**Figure 4:**
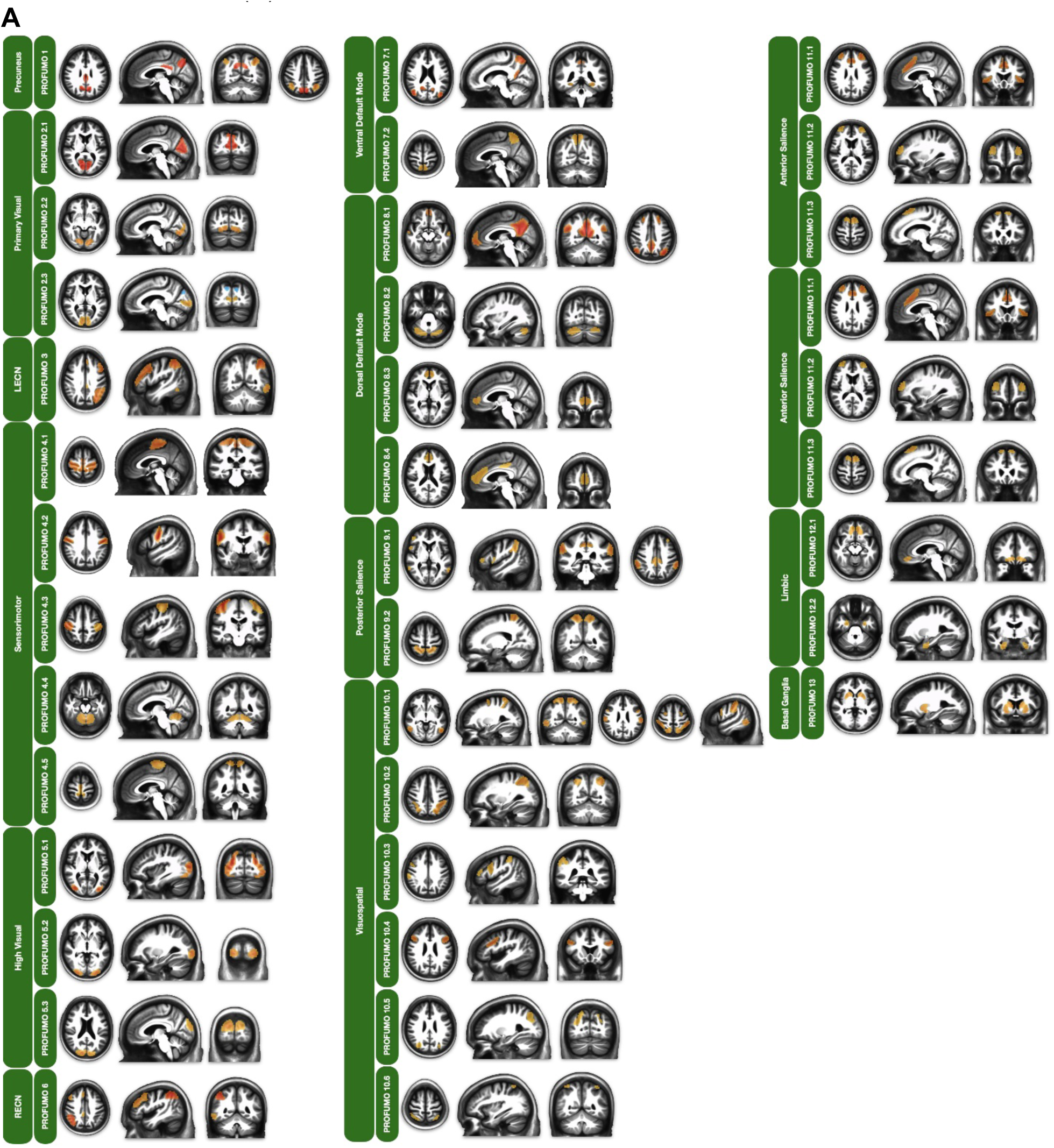
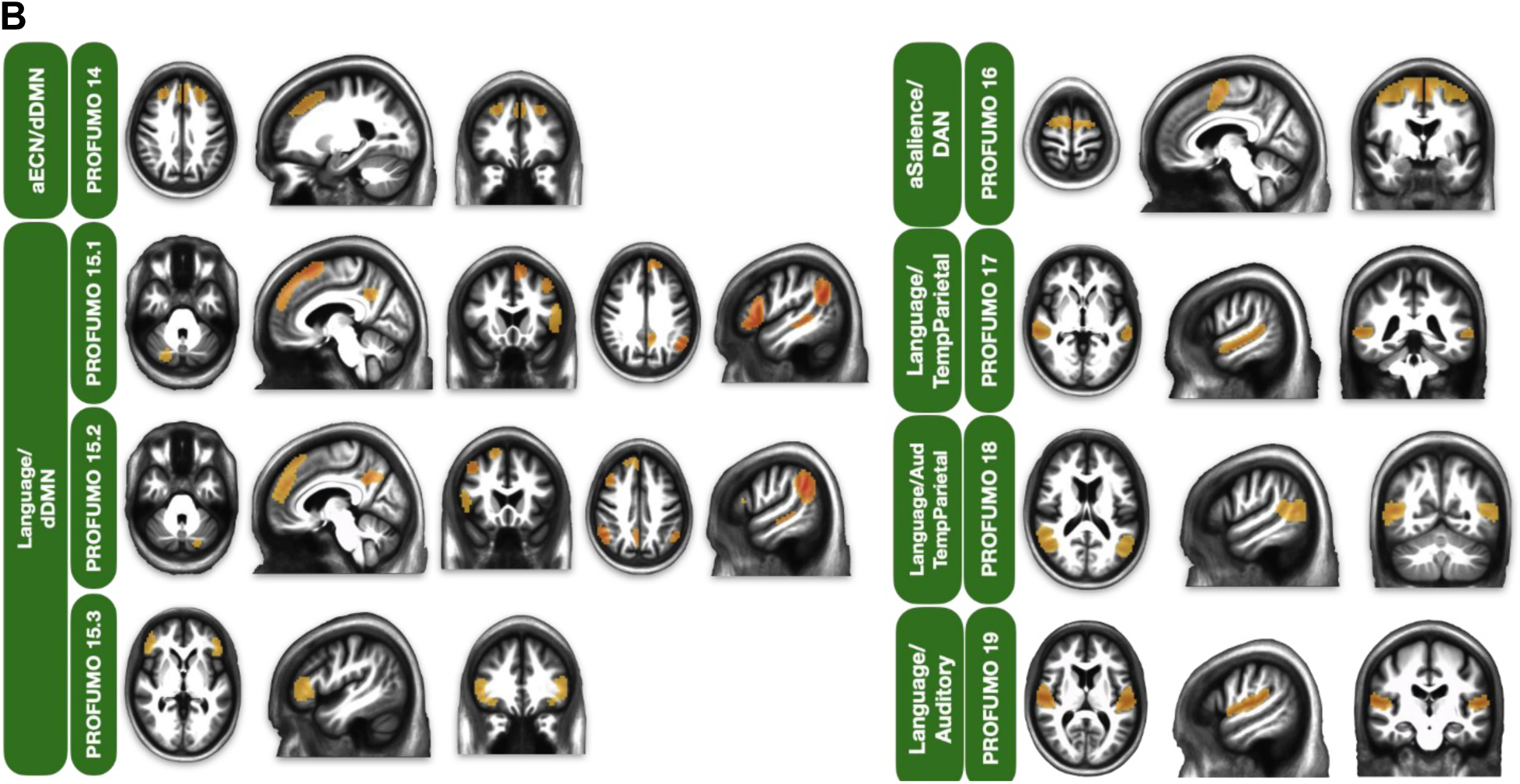
Representative spatial maps of RSN derived from 420 hypertensive older adults with a high familial risk of dementia or subjective cognitive decline, participants in the rrAD trial. Probabilistic Functional Modes (PROFUMO) was performed on resting-state fMRI data using FSL’s newly integrated PROFUMO software. Displayed RSNs are from the 50-mode parcellation, which showed the most ideal network separation from those tested in single networks (A) and combined networks (B).

Three modes represented the primary visual network, kept separate because each captured a distinct temporal configuration, suggesting different temporal dynamics. One mode resembles the GICA map; another is localized to the inferior occipital gyrus, and a third is centered on the middle occipital gyrus, with anticorrelated regions in the superior occipital gyrus.

The sensorimotor network (SMN), consolidated into two components in GICA, was separated into five distinct configurations in PROFUMO. These configurations included the superior, middle, and inferior sections of the pre-/postcentral gyri, along with cerebellar regions identified as independent modes.

The DMN components were further subdivided in PROFUMO. The vDMN, represented as a single component in GICA, was divided into two configurations: one consistent with the GICA map and another focusing on a specific precuneus region. Similarly, the dDMN, the largest RSN in GICA, was separated into four configurations in PROFUMO: canonical regions; cerebellar components, namely the right and left posterior cerebellum; a bilateral prefrontal connection; and the anterior/middle cingulate cortex.

For the salience networks, the posterior salience network was split into two configurations. One included broader regions such as the supramarginal gyrus and frontal areas, while the other was restricted to bilateral superior parietal lobules. The anterior salience network was divided into three configurations, reflecting variations in regional involvement. The visuospatial network, which GICA represented as a combination of the dorsal attention network (DAN) and visuospatial regions, was identified across six separate configurations in PROFUMO. These configurations provided distinct representations of parietal, frontal, and intraparietal sulcus regions. The limbic network, which included the temporal poles and pallidum in GICA, was divided into two configurations in PROFUMO. One mode focused on the orbitofrontal cortex, while the other represented the temporal pole.

For our atlas, two modes of the rrAD420 PROFUMO parcellation resembled configurations of the limbic network. First, we identified a limbic configuration that contained the orbitofrontal cortex. Our second limbic configuration included the temporal pole. A single mode encompassed the entire basal ganglia network.

However, PROFUMO highlighted additional temporal dynamics not apparent in GICA. In networks, such as the high visual network, GICA identified a single bilateral prestriate cortex connection. In contrast, PROFUMO delineated three configurations involving the prestriate cortex, inferior occipital gyrus, and superior occipital gyrus. The auditory network, consistent across methods, showed no significant structural differences.

In addition to the primary "single" networks, the PROFUMO portion of the rrAD420 functional atlas revealed six combinatory networks arising from multimodal regions spanning multiple modes. One of the most prominent combined networks identified was the anterior ECN (aECN)/dDMN network, which encompassed the superior frontal sulcus and medial superior frontal gyrus. Three distinct combinatory configurations involving the language network and dDMN were identified, each providing unique spatial patterns. The first configuration integrated regions such as the right angular gyrus, left angular gyrus, right middle temporal gyrus, left posterior cerebellum, right pars triangularis, right middle frontal gyrus, right dorsomedial prefrontal cortex, and right precuneus. The second configuration involved the left angular gyrus, left middle temporal gyrus, left posterior cerebellum, left pars triangularis, left middle frontal gyrus, left dorsomedial prefrontal cortex, and left precuneus. The final configuration presented a single connection between the right and left pars triangularis.

Another combinatory network observed was the aSalience/DAN network, which comprised two regions in the right and left superior precentral gyri. Additionally, the PROFUMO analysis identified three combinatory networks involving the language network with other functional domains. The first combined the language network with the temporoparietal network, as described by Yeo and colleagues (Thomas Yeo et al. 2011), encompassing the right and left middle temporal gyrus. The second combinatory network spanned the language, auditory, and temporoparietal networks, including regions in the right and left posterior middle temporal gyri and portions of the posterior superior temporal gyrus. The final network represented an overlap between the language network and auditory network, focused within the superior temporal gyrus.

## Discussion

Functional connectivity is a dynamic and evolving process shaped by a complex interplay of intrinsic biological factors and extrinsic influences, such as aging and life experiences (Battaglia et al. 2020). As the brain ages, distinct neuroanatomical and functional changes emerge, including cortical thinning, altered network dynamics, and disruptions in global connectivity. These changes highlight the limitations of functional atlases derived from younger adults, which may fail to capture the unique patterns of connectivity, atrophy, and compensation mechanisms that characterize older populations (Doucet et al. 2021; Zhu et al. 2010).

The rrAD trial’s large cohort of older adults provides a unique opportunity to develop a tailored anatomical template and functional atlas specifically for this age group. Harmonized rs-fMRI scan protocols across scanners enabled the consistent identification of RSNs within this standard space. The rrAD420 template and functional atlas integrate multimodal neuroimaging data (structural and functional) from an older population into a common space, accounting for cohort-specific distinctions, such as cortical atrophy and enlarged ventricles. It complements existing parcellations by introducing a tailored reference that captures the intricacies of late-life neurobiology, offering a resource for studying the functional and structural architecture of the aging brain.

As the prevalence of age-related neurodegenerative diseases rises, it becomes increasingly important to develop functional atlases that accurately reflect late-adulthood connectivity patterns. The rrAD420 functional atlas provides multiple spatial maps, enabling researchers to tailor their analyses to specific study objectives. Crucially, this atlas includes representations of major RSNs such as the DMN, salience network, ECN, sensorimotor network, and primary visual network, all of which have been implicated as key biomarkers of neurodegenerative diseases, including Alzheimer’s disease (Hohenfeld et al. 2018). Additionally, it incorporates less frequently explored but equally important networks, such as the visuospatial, basal ganglia, and limbic networks, broadening the scope for future biomarker discovery and mechanistic investigations in aging research.

The DMN has received considerable attention in the context of aging and neurodegenerative disease due to its selective vulnerability (Andrews-Hanna et al. 2007; Buckner et al. 2008; Damoiseaux et al. 2008; Greicius et al. 2004). It is a task-negative resting-state network composed of interconnected regions like the prefrontal cortex, precuneus, and posterior cingulate cortex (Hohenfeld et al. 2018; Raichle et al. 2001). It plays a critical role in memory and internally driven cognitive functions such as mind-wandering and preparation for future tasks (Raichle 2015), and its dysfunction has been directly linked to cognitive decline in Alzheimer’s disease (AD) (Buckner et al. 2008).

Disrupted DMN connectivity is a well-established hallmark of AD. For example, lower DMN connectivity has been observed in individuals with mild cognitive impairment (MCI) who subsequently progress to AD, suggesting its potential as a biomarker for disease progression (Van Dijk and Sperling 2011; Zhu et al. 2013). This dysfunction is further supported by studies showing a link between reduced DMN connectivity and cognitive impairments in prodromal AD (Koch et al. 2015). Importantly, AD hallmarks such as amyloid-beta (Aβ) and hyperphosphorylated tau accumulate in key DMN regions, a process potentially linked to sustained high activity of DMN neurons (Putcha et al. 2022; Šimić et al. 2014; Sperling et al. 2009). Our own recent research supports this, showing a direct correlation between the amplitude of BOLD fluctuations and Aβ deposition, particularly in DMN regions (Scheel et al. 2021). The inclusion of multiple DMN-specific atlases in our rrAD420 functional atlas provides a valuable resource for precisely examining how changes in DMN structure and function are linked to age-related disease processes in this at-risk population.

Beyond its utility for the rrAD cohort, the rrAD420 template and its associated flow fields have broader translational potential, providing a more representative foundation for studying RSNs across diverse older populations. By integrating high-quality baseline data from the rrAD study and harmonizing images across multiple modalities, this atlas enhances anatomical and functional accuracy. This makes the rrAD420 functional atlas an invaluable tool for researchers seeking to unravel the complexities of brain aging, providing a resource that bridges the gap between cohort-specific characteristics and universal principles of late-life neural organization.

### Seed-based Correlation Analysis

The rrAD420 atlas includes a high-resolution, seed-based spatial map of DMN (DMN24) tailored for older adults. It will benefit primarily researchers concerned with the DMN and its potential age-related alterations. The inclusion of the DMN24, which encompasses six cerebellar regions, is particularly significant, as recent studies have highlighted the cerebellum’s role in DMN connectivity (Guell et al. 2018). The cerebellum is often overlooked in functional neuroimaging because it is excluded from the field of view due to head position or brain size (Wang et al. 2025). Our sample provided sufficient cerebellar coverage to estimate functional connectivity at these locations, and the resulting connectivity regions were supported by the existing literature (Buckner et al. 2008). When the cerebellum is excluded, the remaining DMN regions form a subnetwork of DMN24, referred to as DMN18. Although these six regions showed strong functional connectivity, providing both the DMN24 and DMN18 spatial maps accommodates a broader range of research objectives, allowing investigators to either leverage the comprehensive network, including the cerebellum, or focus solely on cortical and subcortical regions.

### Group Independent Component Analysis

The GICA portion of the rrAD420 functional atlas provides spatial maps for all 15 RSNs identified across the whole brain. This method represents the current standard for data-driven RSN identification and will help investigate independent RSNs in older populations. The GICA parcellation demonstrates several unique and valuable features compared to existing RSN atlases (Damoiseaux 2017), providing deeper insights into functional connectivity in older adults with a high familial risk of dementia. Such examples include characterizing the precuneus network as separate from the DMN. While the precuneus is typically regarded as a core region of the DMN, its segregation in this cohort, along with the inclusion of posterior/middle cingulate cortex regions, suggests unique functional specialization. This separation highlights potential alterations in connectivity patterns associated with aging, hypertension, or early cognitive decline.

The richness of the dorsal DMN (dDMN) in the rrAD420 atlas is particularly interesting. Two components of our GICA parcellation resembled the dDMN. These components were combined to create a single dDMN spatial map. The final dDMN spatial map contained the most brain regions of any RSN in the rrAD420 functional atlas. This reflects its multifaceted role in self-referential processing, episodic memory, and higher-order cognition (Menon 2023). The detailed parcellation of the dDMN underscores its importance in supporting complex cognitive functions and its potential as a marker for early disruptions in neurodegenerative conditions.

Another intriguing aspect is the differentiation between the RECN and LECN, with their distinct functional roles and potentially distinct temporal dynamics despite their regional overlap. The RECN appears more involved in externally directed attention and task control, whereas the LECN demonstrates stronger ties to internally directed cognitive processes (Kazemi-Harikandei et al. 2022). This differentiation provides valuable insights into how these networks might interact or compete in the context of cognitive aging and disease.

The atlas also shows the bilateral connections of the auditory network. The high visual network regions of the lateral occipital lobe consisted of a single bilateral connection between the right and left prestriate cortices. The DAN/Vis network contained a mixture of regions from the visuospatial network, as presented in the Shirer atlas, as well as regions that belong to the DAN, presented by Yeo and colleagues (Shirer et al. 2012; Yeo et al. 2011). Specifically, the DAN/Vis network included the superior parietal cortex, precuneus, superior frontal cortex, inferior frontal gyrus, middle temporal gyrus, and right inferior temporal gyrus. The atlas resolves some small spatial conflicts through manual cleanup in the salience networks, where the posterior salience network comprises the supramarginal gyrus, precuneus/middle cingulate cortex, thalamus, insula, and superior cerebellum. Regions of the anterior salience network were identified across four components and were subsequently combined into a single spatial map to capture the RSN in its entirety. The final anterior salience network map included the superior frontal gyrus, middle frontal gyrus, dorsal anterior cingulate cortex, superior frontal gyrus/pre-supplemental motor area, insula, superior cerebellum, and inferior cerebellum. We also observed the language network across two components, which were merged to create a finalized spatial map of the language network.

In sum, the GICA parcellation is intended to help studies investigating changes in functional connectivity across entire RSNs of interest. However, this technique presents certain drawbacks, including its unidirectional nature, which does not explicitly account for inter-individual variability, and its limited capacity to differentiate between overlapping RSNs.

### Probabilistic Functional Modes

The PROFUMO portion of the rrAD420 functional atlas extends GICA by simultaneously modeling RSNs at both group and participant levels via Bayesian inference (Harrison et al. 2015). This dual-level modeling enabled the PROFUMO approach to delineate a greater proportion of modes derived from true signal origins, reducing the influence of noise and minimizing the exclusion of meaningful data. Consequently, the PROFUMO parcellation retains a comprehensive representation of functional connectivity, providing researchers with valuable insights into network-specific characteristics and variability.

PROFUMO also enabled grouping multimodal regions into various configurations and combinatory networks, facilitating a deeper exploration of RSN subdivisions. These factors offer advantages over GICA in some cases, as they accommodate the study of specific RSN subdivisions that can vary in their susceptibility to connectivity disruptions in age-related diseases (Viviano et al. 2019; Xue et al. 2019). Namely, the primary visual, SMN, high visual, vDMN, dDMN, posterior salience, visuospatial, anterior salience, and limbic networks of the PROFUMO parcellation were represented by multiple configurations. The configuration that best captured the primary representation of each RSN was listed first for each respective network. While the primary configurations align closely with those derived from GICA, they are not identical, underscoring PROFUMO’s ability to reveal network-specific variations. This distinction highlights the utility of PROFUMO for applications that require precise delineation of RSN sub-configurations, particularly for studying age-related alterations or disease-specific connectivity disruptions. However, other networks, such as the Basal Ganglia, appear as singular modes, as expected given the relatively unified nature of this network (Wen et al. 2012).

An important contribution of the PROFUMO atlas lies in its identification of overlapping RSNs that were highly correlated with one another and of combinatory networks. For instance, the DMN has been shown to have interconnections with the language network (Gordon et al. 2020). The DMN recruitment of language network regions, such as the middle temporal gyrus and middle frontal gyrus, aligns with existing evidence that these regions are recruited during processes like inner speech or verbal thinking (Alderson-Day et al. 2016). These combinations underscore the shared functional roles of the language and default mode regions in semantic processing, memory retrieval, and complex cognitive integration. The inclusion of the cerebellum emphasizes its involvement in higher-order cognitive processes, such as language and attention. The functional overlap between the dDMN and language networks in the PROFUMO parcellation provides an opportunity to explore biomarkers of cognitive function and their disruption in neurodegenerative diseases.

Additional combinations that align with existing evidence: The aECN/ dDMN combination reflects the integration of executive control, self-referential processing, and goal-directed behavior (Liang et al. 2016). It highlights the intricate relationship between cognitive control mechanisms and default mode functions, such as introspection and planning, which are critical for flexible and adaptive cognition. The combination aSalience/DAN points to integration of attentional control with salience detection in regions traditionally associated with motor planning and execution. The convergence of salience and attentional networks in these areas suggests a role in prioritizing goal-relevant stimuli during motor-related tasks. The combinatory networks, spanning the language network, auditory network, and temporoparietal network, demonstrate phonological and semantic integration critical for integrating auditory and linguistic information, facilitating communication and comprehension (Turken and Dronkers 2011). In sum, the combinatory networks highlight the added value of the PROFUMO parcellation by enabling the identification of multimodal interactions and functional overlaps across RSNs.

By allowing regions to belong to multiple modes, PROFUMO reveals the complex and dynamic interplay between networks and captures functional heterogeneity consistent with recent advances in network neuroscience. Across all parcellation approaches, the rrAD420 atlas effectively captures the cerebellum’s emerging role in higher-order cognitive functions by including its regions within multiple combinatory networks. This is supported by recent research that implicates the cerebellum in cognitive control and attentional processes (D’Mello et al. 2020). These findings underscore the rrAD420 atlas’s unique ability to delineate complex network relationships and provide a foundation for future studies exploring the cerebellum’s contributions to functional connectivity.

### rrAD420 – multiple modalities

The rrAD420 atlas represents a significant step forward in multimodal neuroimaging research, offering a comprehensive framework for studying both normal aging and pathological changes. Beyond rs-fMRI data, the rrAD study incorporated additional imaging modalities, including T2-FLAIR for quantification of white matter hyperintensity (WMH), arterial spin labeling (ASL) for estimating regional cerebral blood flow (CBF), and diffusion imaging for assessing fiber tracts and structural connectivity. By aligning all data to the rrAD420 space, the atlas enables integrative analyses across multiple imaging modalities, providing a unique opportunity to examine the interplay among structural, functional, and vascular changes in the aging brain.

### Limitations

While this study utilized state-of-the-art neuroimaging techniques to develop an age-appropriate structural template and functional atlas, several limitations should be acknowledged. First, the rrAD cohort consists exclusively of hypertensive older adults, which may restrict the generalizability of the atlas to non-hypertensive populations. Hypertension is known to influence brain structure and function, including alterations in functional connectivity and white matter integrity, which could introduce biases specific to this population (Hajjar et al. 2016; Iadecola and Davisson 2008). However, given the high prevalence of hypertension and other CVD in older adults, with rates exceeding 75% in the 60-79 age group, the rrAD420 atlas remains a valuable resource for studies involving this demographic. It provides a more representative alternative compared to atlases derived from younger or mixed-age cohorts.

## Conclusion

The rrAD420 atlas represents a significant contribution in neuroimaging for aging populations. Unlike widely used atlases, which are based on younger, healthy adults, the rrAD420 atlas is specifically tailored for older adults and accounts for age-related structural and functional changes that are often overlooked in general-purpose atlases. By incorporating imaging data from 420 older adults with hypertension from the rrAD clinical trial, we developed a comprehensive suite of functional atlases on a common template tailored to older populations. Using a hypothesis-driven seed-based approach, we constructed two DMN-specific atlases: DMN24, which includes cerebellar regions, and DMN18, which excludes them. Additionally, we applied a data-driven GICA approach to create a functional atlas termed "rrAD420-GICA," characterized by non-overlapping spatial maps, and the Bayesian PROFUMO approach to develop "rrAD420-PROFUMO," which accommodates spatial overlap and identifies multimodal and combinatory RSNs. These distinct atlases offer unique strengths for studying functional connectivity. The GICA-based atlas is ideal for global network analyses with its non-overlapping spatial maps, while the PROFUMO-based atlas captures more complex, overlapping configurations and combinatory networks.

This versatile, multimodal platform is not only instrumental for analyzing the rrAD trial outcomes but also serves as a critical resource for broader research into aging and neurodegenerative diseases. By integrating various imaging data within a unified space (e.g., white matter hyperintensities, cerebral blood flow, diffusion metrics), the rrAD420 atlas supports the identification of robust biomarkers. It provides a crucial foundation for future research aimed at aiding in the prognosis, diagnosis, and treatment of these conditions.

## Acknowledgements

This work has been supported by NIH grants (R01AG049749, R01AG057571, R56AG074613 and RF1AG084134). The high-performance computational resources were provided by the Institute for Cyber-Enabled Research at Michigan State University.

## Data Availability Statement

The rrAD420 Toolbox, comprising the anatomical templates, functional parcellations (NIfTI format), and associated code, is available for download as a complete, integrated package on Zenodo (doi: https://doi.org/10.5281/zenodo.17727416). The source code is also maintained for public review on GitHub at https://github.com/NOSrepo/NOS_rrAD420, which includes instructions for integrating the data if downloading the code separately.

## Code Availability Statement

The analysis code specifically developed for this study is available in the Zenodo repository listed above. Standard software packages used include SPM12 (fil.ion.ucl.ac.uk/spm), FSL (fsl.fmrib.ox.ac.uk), FreeSurfer (surfer.nmr.mgh.harvard.edu), and AFNI (afni.nimh.nih.gov).

## Author Contributions

**N.S., Z.F.**: Conceptualization, Methodology, Formal analysis, Writing – original draft. **J.B., P.Y.**: Data curation and interpretation, Writing – review & editing. **J.K., E.B., E.V., J.B., A.S., D.K., M.C., L.H., W.V., R.Z.**: Investigation, Resources, Funding acquisition, Writing – review & editing. **D.C.Z.**: Supervision, Conceptualization, Project administration, Writing – review & editing.

